# STRUCTURE-BASED DESIGN OF PROTACS FOR THE DEGRADATION OF SOLUBLE EPOXIDE HYDROLASE

**DOI:** 10.1101/2024.09.05.611393

**Authors:** J. Schönfeld, S. Brunst, L. Ciomirtan, N. Liebisch, A. Kumar, J. H. M. Ehrler, L. Wintermeier, J. Heering, A. Brüggerhoff, L. Weizel, A. Kaiser, M. Schubert-Zsilavecz, S. Knapp, R. Fürst, E. Proschak, K. Hiesinger

**Affiliations:** Institute of Pharmaceutical Chemistry, Goethe University, 60438 Frankfurt/Main, Germany; Department of Pharmacy - Center for Drug Research, Ludwig-Maximilians-University Munich, Butenandtstr. 5-13, 81377 Munich, Germany; Fraunhofer Institute for Translational Medicine and Pharmacology ITMP, Theodor-Stern-Kai 7, 60596 Frankfurt/Main, Germany

## Abstract

Soluble epoxide hydrolase (sEH) represents a promising target for inflammation-related diseases as it hydrolyzes highly anti-inflammatory epoxy-fatty acids (EpFAs) to the less active corresponding diols.^1^ sEH harbours two distinct catalytic domains, the C-terminal hydrolase domain and the N-terminal phosphatase domain which are connected by a proline-rich linker.^2^ Although potent inhibitors of enzymatic activity are available for both domains, sEH-PROTACs offer the unique ability to simultaneously degrade both domains, mimicking the sEH knockout phenotype associated with beneficial effects as reducing inflammation, attenuating neuroinflammation, and delaying the progression of Alzheimer’s disease. Herein, we report the structure-based development of a potent sEH-PROTAC as a useful tool compound for the investigation of sEH. In order to facilitate a rapid testing of the synthesized compounds a cell-based sEH degradation assay was developed based on the HiBiT-technology. A structure-activity-relationship (SAR) investigation was performed, based on the crystal structure of previously published sEH inhibitor FL217 where we identified two possible exit vectors. We designed and synthesized a set of 24 PROTACs with varying linkers in a combinatorial manner. Furthermore, co-crystallization of sEH with two selected PROTACs allowed us to explore the binding mode and rationalize the appropriate linker length. After biological and physicochemical investigation, the most suitable PROTAC **23** was identified and applied to degrade sEH in primary human macrophages, marking the successful translation and applicability to non-artificial systems.

## Introduction

Soluble epoxide hydrolase (sEH) is a ubiquitously expressed bifunctional enzyme encoded by the gene EPHX2. The protein harbours two distinct catalytic domains connected by a proline-rich linker. The C-terminal domain exhibits epoxide hydrolase activity (sEH-H), while the N-terminal domain (sEH-P) represents a lipid phosphatase.^2,3^ sEH plays a pivotal role in the CYP-branch of the arachidonic acid cascade as its epoxide hydrolase domain (sEH-H) converts anti-inflammatory epoxyeicosatrienoic acids (EETs) into the corresponding biologically less active diols.^4,5^ Since sEH metabolism is the main pathway for the deactivation of EETs^6^, it is being pursued as a pharmacological target for multiple inflammatory diseases, and several sEH inhibitors have been developed^7^, e.g. clinical candidate GSK2256294A (**1**)^8–11^ (figure 1). The biological role of sEH-P is less well understood and the endogenous substrates are still being investigated^12–14^.

**Figure 1:**
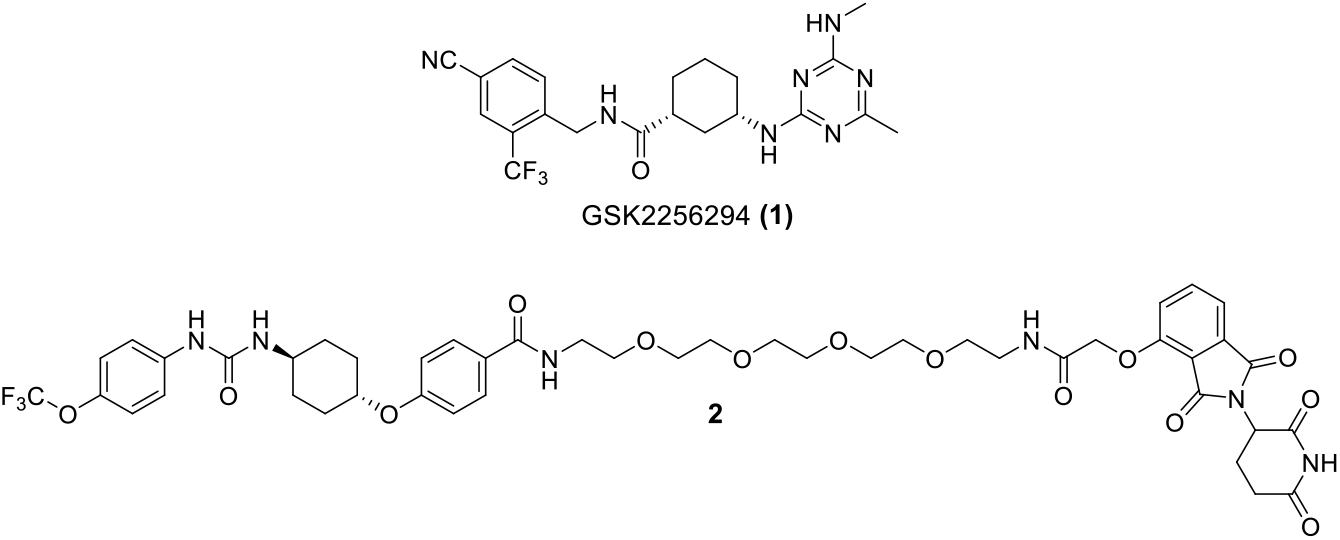
Structures of the potent sEH-H inhibitor and clinical candidate GSK2256294A (**1**) as well as the first-in-class sEH PROTAC **2**

Depletion of the whole enzyme in sEH knockout mouse models is associated with beneficial effects as reducing inflammation,^15^ attenuating neuroinflammation^16^, and delaying the progression of Alzheimer’s disease^17^. Thus, simultaneous inactivation of both domains could be therapeutically advantageous and would also offer the opportunity to further study the biological functions of sEH. Although there are potent inhibitors available for both domains^7,13^, the application of multiple drugs is often associated with negative safety profiles and drug-drug-interactions.^18^ sEH-PROTACs however offer the unique opportunity to simultaneously degrade both domains, mimicking the sEH knockout phenotype. Proteolysis targeting chimeras (PROTACs) are heterobifunctional molecules that induce the ubiquitinylation and subsequent proteasomal degradation of a protein of interest (POI) by bringing the POI into close proximity with an E3 ligase as e.g. cereblon (CRBN) or the Van-Hippel-Lindau E3 ligase (VHL).^19^ Recently, Wang et al. reported the development of a first-in-class series of sEH-PROTACs. Their lead compound **1a** (figure 1) induced sEH degradation in HepG2 and HEK293T cells. Remarkably, **1a** was also more effective in reducing ER stress in a phenotypic cellular assay compared to the parent sEH inhibitor.^20^ Peyman et al. likewise demonstrated an attenuation of ER stress in mice in vivo.^21^

In this study, we developed a cellular sEH degradation assay based on the HiBiT-technology^22^ in order to accelerate the degradation ability of synthesized sEH PROTACs. We performed a structure-activity-relationship (SAR) investigation based on the crystal structure of previously published potent sEH inhibitor FL217(**3**).^23^ To demonstrate the applicability to study sEH in cellular context, the most promising PROTAC was studied in primary human cells.

## Results and Discussion

### Design strategy

Our rational design approach to develop sEH PROTACs was based on the previously published crystal structure of the potent sEH inhibitor FL217 **(3)** (IC_50_ (hsEH) = 8.4 nM *in vitro*) shown in figure 2.^23^ The hydrolase domain of sEH (sEH-H) possesses a hydrophobic L-shaped binding pocket exhibiting a long branch (**∼**15 Å) and a short branch (**∼**10 Å).^23,24^ Amide-based inhibitor **3** functions as a transition state mimetic of the endogenous epoxide substrates: Tyr383 and Tyr466 residues in the active site are coordinating to the carbonyl-O via two H-bonds, while Asp335 interacts with the amide-N. Analyzing the crystal structure, we identified two possible exit vectors to address either the exit of the short branch or the long branch of the binding pocket. We rationalized that replacing the cyclopropane ring and attaching a linker to the sulfonamide moiety of **3** would lead to a short branch addressing inhibitory scaffold, whereas functionalization at the Indole-N would result in a long branch addressing scaffold (figure 2). Molecular docking experiments were performed in order to identify the appropriate linker length for the inhibitory scaffold to reach the exit of the binding pocket (figure S1). Based on these results, we decided to modify **3** at both attachment points with a C3-alkyl linker bearing a terminal alkyne group and obtained short branch addressing scaffold **4** and long branch addressing scaffold **5** (figure 2). The terminal alkyne handle allowed for an efficient variation of linkers and E3 ligase recruiters using the Cu(I) catalysed azide-alkyne cycloaddition (CuAAC, “CLICK” reaction) as well as the Sonogashira coupling reaction.

**Figure 2:**
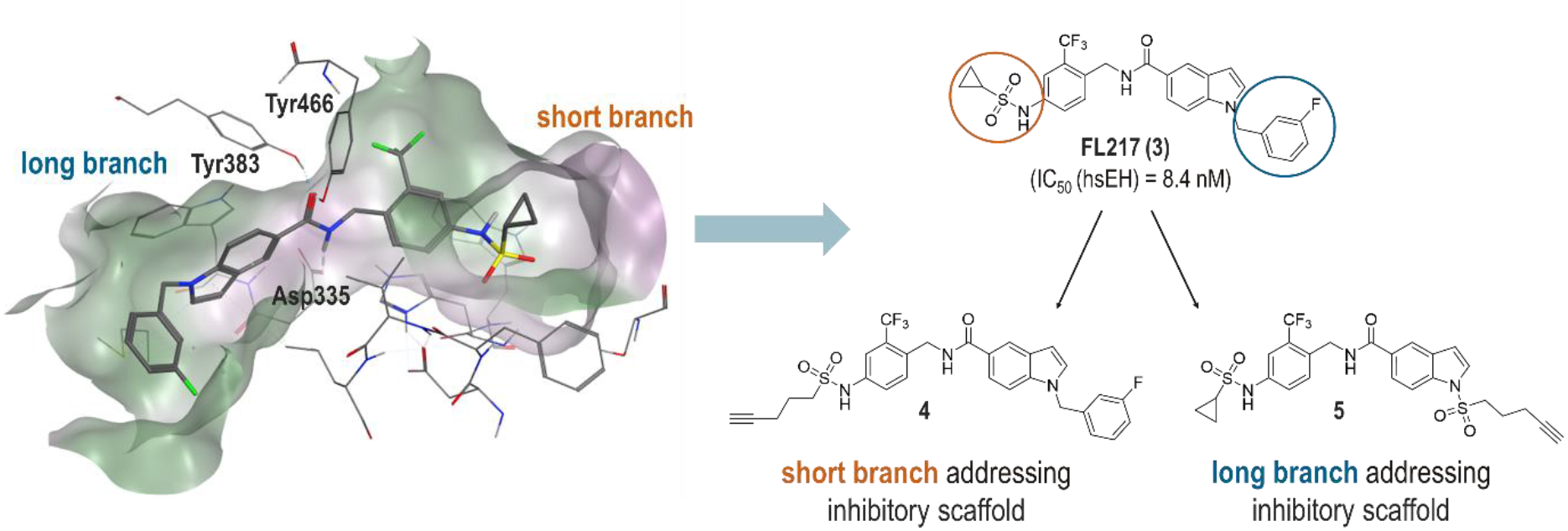
Structure-based design of sEH PROTACs. Left: Crystal structure of sEH, co-crystallized with sEH inhibitor FL217 (**3**) (PDB: 7P4K). Right: Two inhibitory scaffolds were developed from sEH inhibitor FL217, each bearing a terminal alkyne group for further functionalization.

In this study, we aimed to recruit the E3 Ligase cereblon (CRBN) and implicated derivatives of the well-established CRBN-ligands thalidomide, pomaliomide, and lenalidomide in our PROTAC design. Linker variation was implemented by the choice of rigid alkyne linkers as well as flexible PEG linkers of different lengths (PEG1-PEG6).

### Synthesis

Short branch addressing inhibitor scaffold **3** was synthesized from aniline precursor **11** in a nucleophilic substitution reaction with pent-4-yne-1-sulfonyl chloride and pyridine. The synthesis of precursor **11** was performed in three steps according to the previously published procedure by Lillich et al.:^23^ Indole ester **6** was substituted with benzyl chloride **7** in a Finkelstein type nucleophilic substitution reaction. After ester hydrolysis, carboxylic acid **9** was treated with benzylic amine **10** under amide coupling conditions to obtain **11** (scheme 1).

**Scheme 1:**
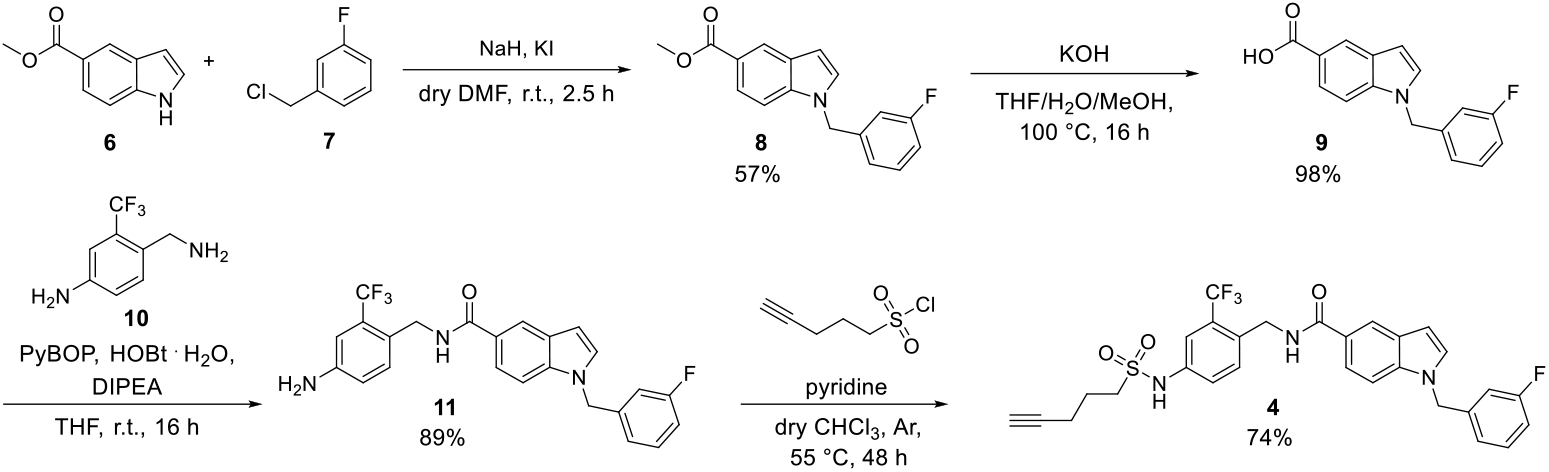
Synthetic route for inhibitor scaffold 4 addressing the short branch of the binding pocket.

The synthesis of long branch addressing scaffold **5** was conducted in four steps. According to the previously published method,^23^ aniline precursor **16** was synthesized from starting material 4-amino-2-(trifluoromethyl)benzonitrile (**12**) in three steps. **12** underwent nucleophilic substitution with cyclopropanesulfonyl chloride, followed by reduction of **13** to benzylic amine **14** and subsequent amide coupling with *1H*-indole-5-carboxylic acid (**15**) to obtain **16**. Lastly, **5** was synthesized from precursor **16** in a nucleophilic substitution reaction with pent-4-yne-1-sulfonyl chloride and NaH (scheme 2).

**Scheme 2:**
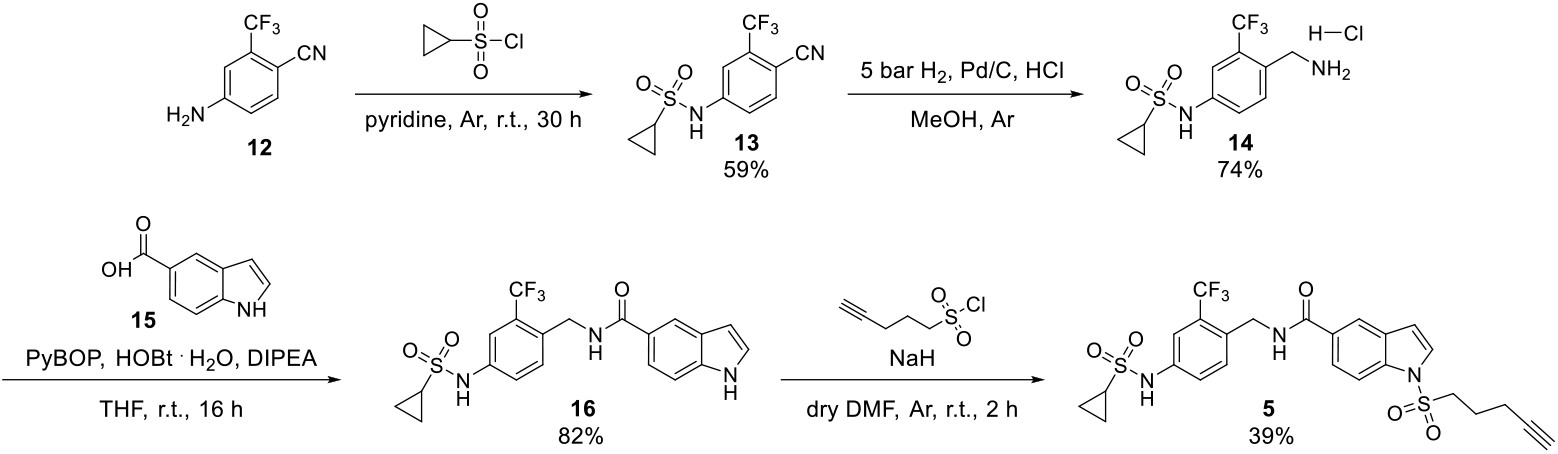
Synthetic route for inhibitor scaffold 5 addressing the long branch of the binding pocket.

The set of PROTACs with rigid alkyne linkers (**18a-e** and **19a-e**) was synthesized under Sonogashira coupling conditions^25^ with both inhibitory scaffolds **4** and **5** (scheme 3). Five commercially available Br-functionalized CRBN recruiters were used from which two were pomalidomide derivatives **17a-17b** (functionalized in position 4 and 5, respectively) and three were lenalidomide derivatives **17c-17e** (Br in position 4, 5 and 6, respectively). With this set of regioisomeric compounds we aimed to examine the possible impact of the attachment point to the CRBN recruiter on sEH degradation. Position 7 was not taken into consideration for this study, as Fischer et al. demonstrated this position not to be solvent exposed in the binding pocket of CRBN.^26^

**Scheme 3.**
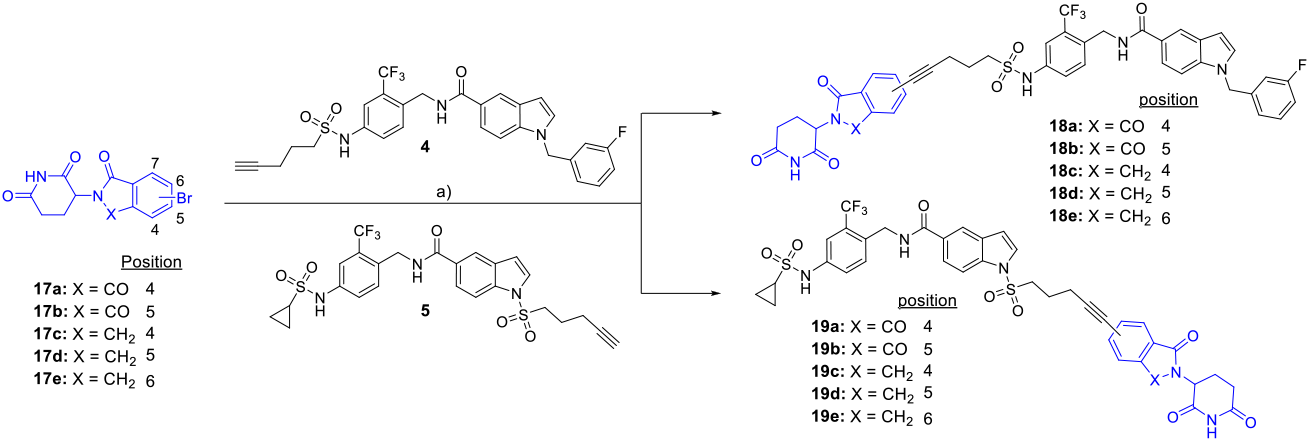
Preparation of PROTACs with alkyne linkers. Reaction conditions (a): Pd(PPh_3_)_2_Cl_2_ (0.1 equiv.), CuI (0.2 equiv.), DMF, NEt_3_, 80 °C, 16 h.

**Scheme 4:**
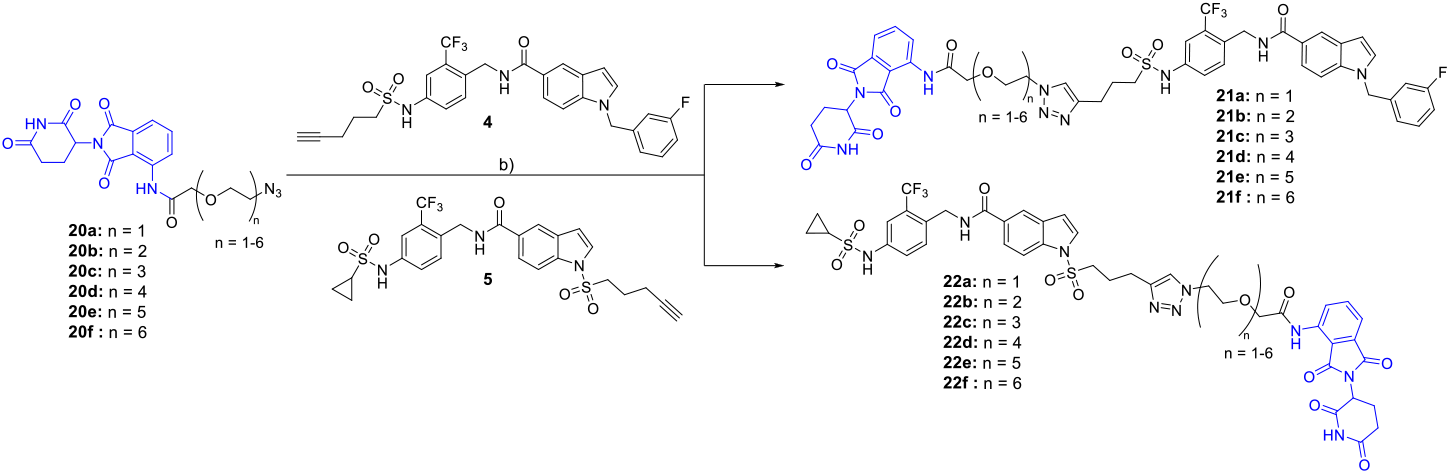
Preparation of PROTACs with PEG linkers. Reaction conditions (b): CuSO_4_. 5 H_2_O (0.3 equiv.), sodium ascorbate (0.3 equiv.), DMF/H_2_O = 4:1, r.t., 16 h.

For the set of PROTACs harbouring PEG linkers, we used commercially available azide functionalized pomalidomide derivatives (**20a-20f**) with linker lengths varying from PEG1 to PEG6. Both inhibitory scaffolds **4** and **5** were treated with **20a-20f** under standard CuAAC conditions^27^ to obtain the corresponding PROTACs **21a-21f** and **22a-22f** (scheme 4).

### Crystal structure

To validate the postulated binding modes of short branch vs. long branch addressing PROTACs, the two PEG2 exhibiting PROTACs **21b** and **22b** were co-crystallized with human sEH (figure 3). The obtained cocrystal structures confirmed our hypothesis: While **21b** (depicted in grey) addresses the exit of the short branch of the binding pocket, **22b** (shown in orange) addresses the short branch exit.

**Figure 3:**
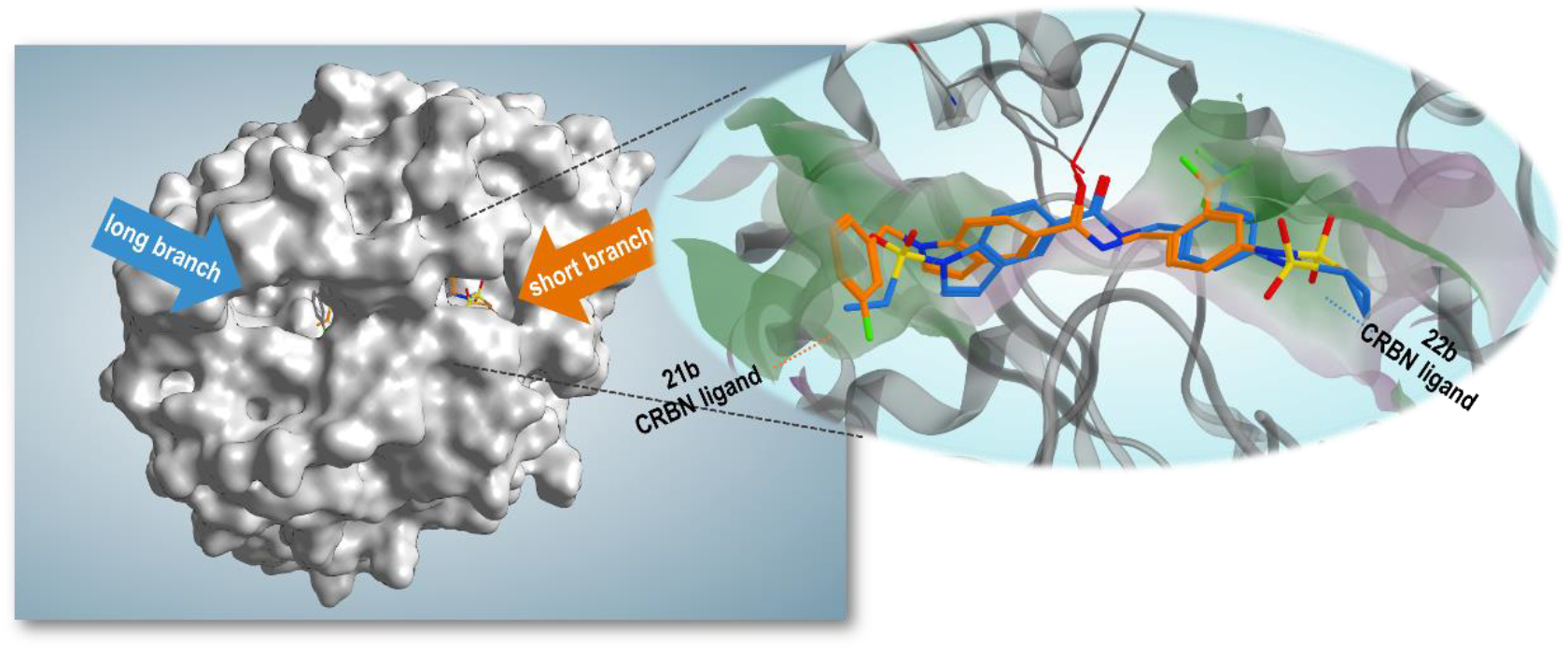
Overlaid cocrystal X-ray structures of and **21b** (orange) and **22b** (blue) with human sEH (PDB codes: 8S76, 8S77). As predicted, **21b** addresses the exit of the short branch of the binding pocket, whereas **22b** addresses the long branch exit. PEG2-linkers and CRBN ligands are not resolved.

### Biochemistry

For all synthesized PROTACs, the in vitro sEH inhibitory potency was determined in an enzyme activity assay with recombinant sEH (murine and human isoforms) and the fluorogenic substrate 3-phenyl-cyano(6-methoxy-2-naphthalenyl)methyl ester-2-oxiraneacetic acid (PHOME).^28^ All the compounds exhibited low nanomolar potencies towards human sEH-H (hsEH-H), confirming that the attached linkers were tolerated in the active site of hsEH-H. Regarding the murine isoform, up to three magnitudes lower potencies were observed for the short branch addressing compounds **18a-e** and **21a-f** (tables 1 and 2) still exhibiting decent activity.

**Table 1:**
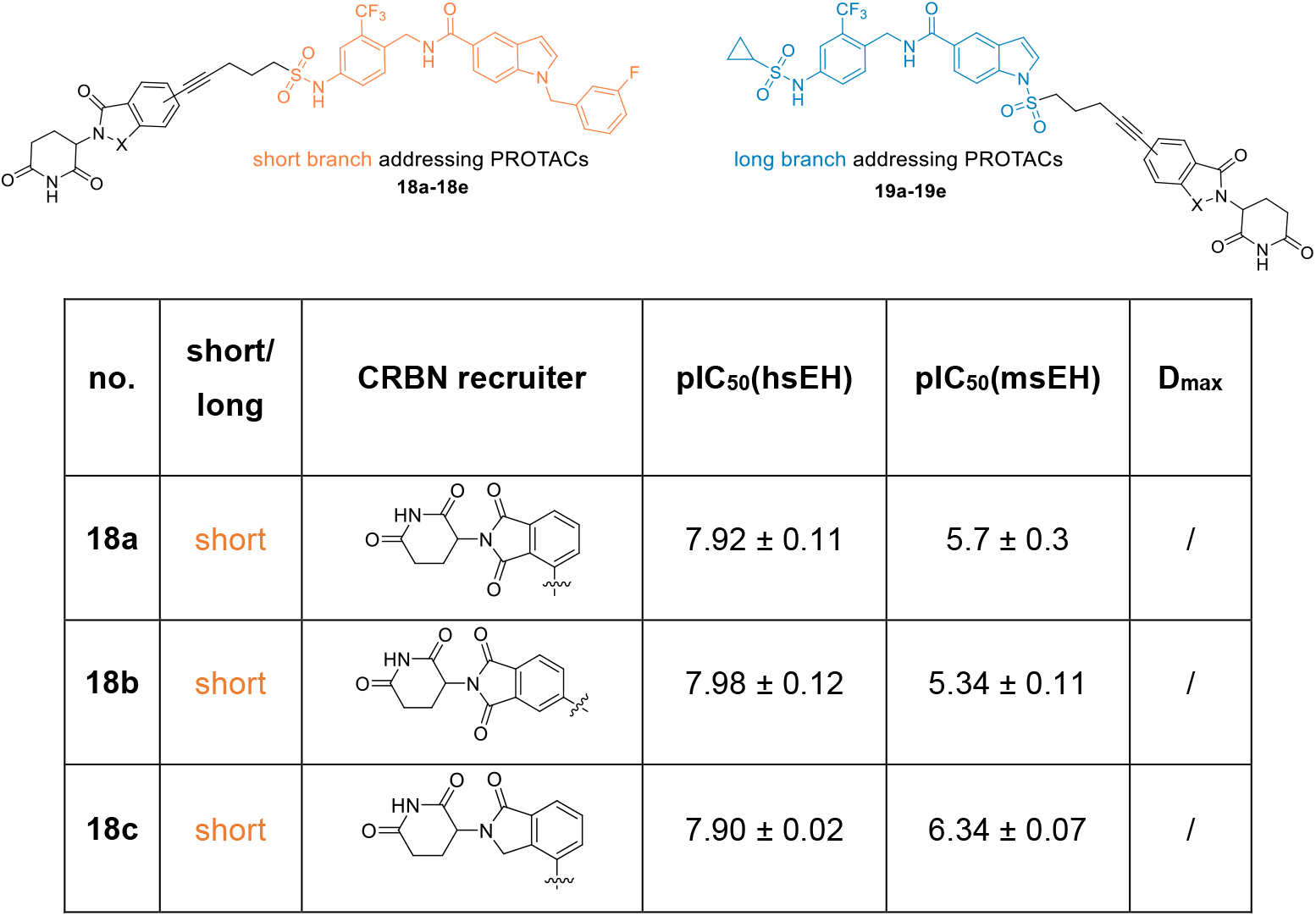
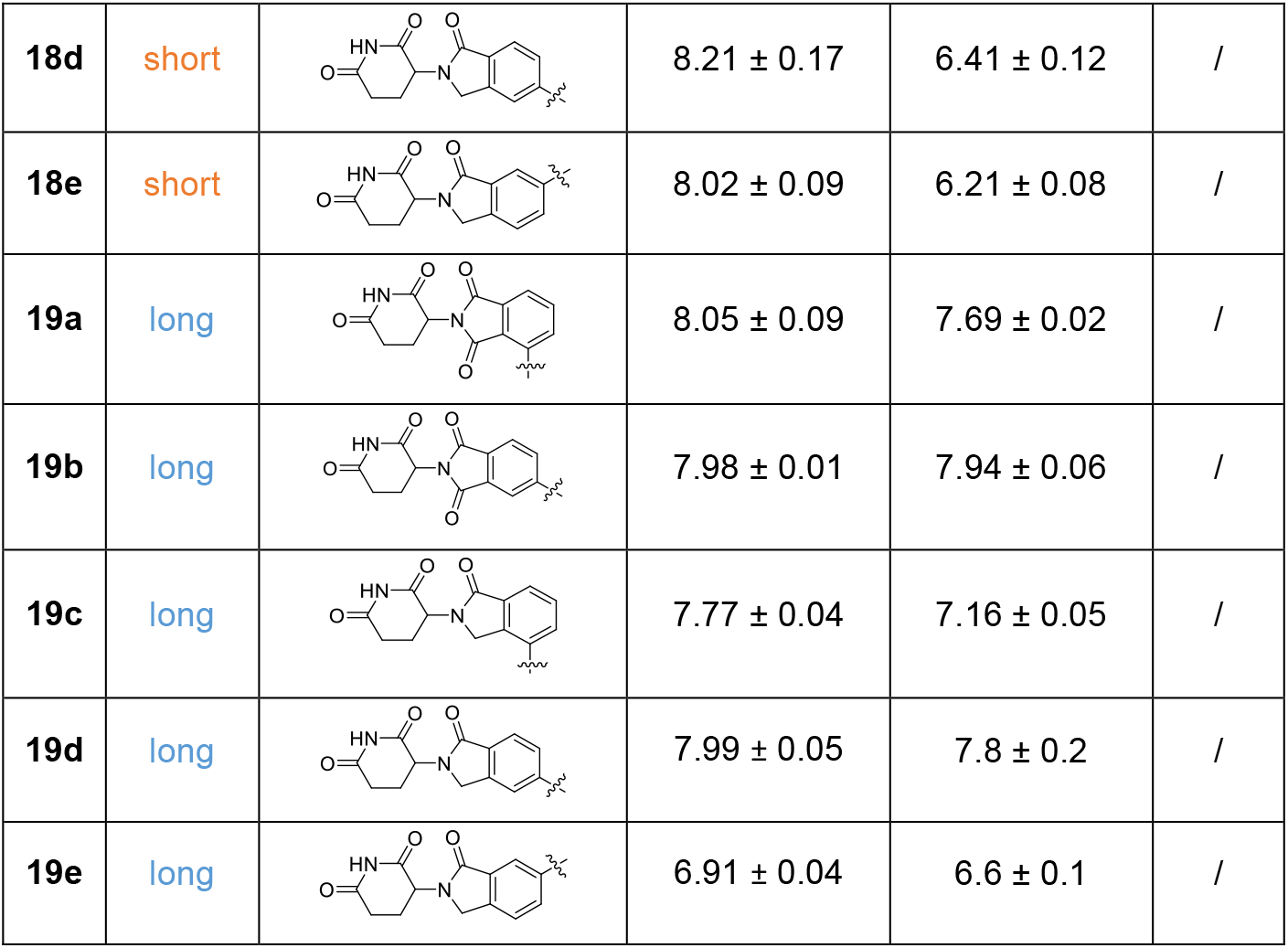
Potential sEH PROTACs with alkyne linkers linked from different positions of the CRBN recruiters.

| no. | short/<br>long | CRBN recruiter | pIC<sub>50</sub>(hsEH) | pIC<sub>50</sub>(msEH) | D<sub>max</sub> |
| --- | --- | --- | --- | --- | --- |
| **18a** | short |  | 7.92 ± 0.11 | 5.7 ± 0.3 | / |
| **18b** | short |  | 7.98 ± 0.12 | 5.34 ± 0.11 | / |
| **18c** | short |  | 7.90 ± 0.02 | 6.34 ± 0.07 | / |
| **18d** | short |  | 8.21 ± 0.17 | 6.41 ± 0.12 | / |
| **18e** | short |  | 8.02 ± 0.09 | 6.21 ± 0.08 | / |
| **19a** | long |  | 8.05 ± 0.09 | 7.69 ± 0.02 | / |
| **19b** | long |  | 7.98 ± 0.01 | 7.94 ± 0.06 | / |
| **19c** | long |  | 7.77 ± 0.04 | 7.16 ± 0.05 | / |
| **19d** | long |  | 7.99 ± 0.05 | 7.8 ± 0.2 | / |
| **19e** | long |  | 6.91 ± 0.04 | 6.6 ± 0.1 | / |

**Table 2:**
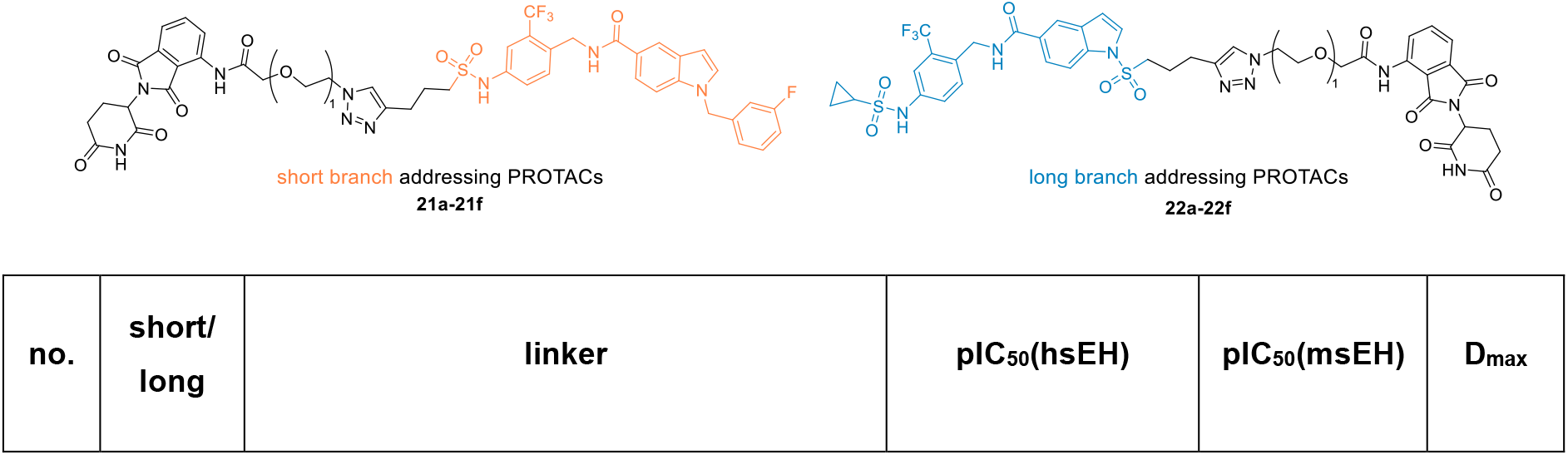
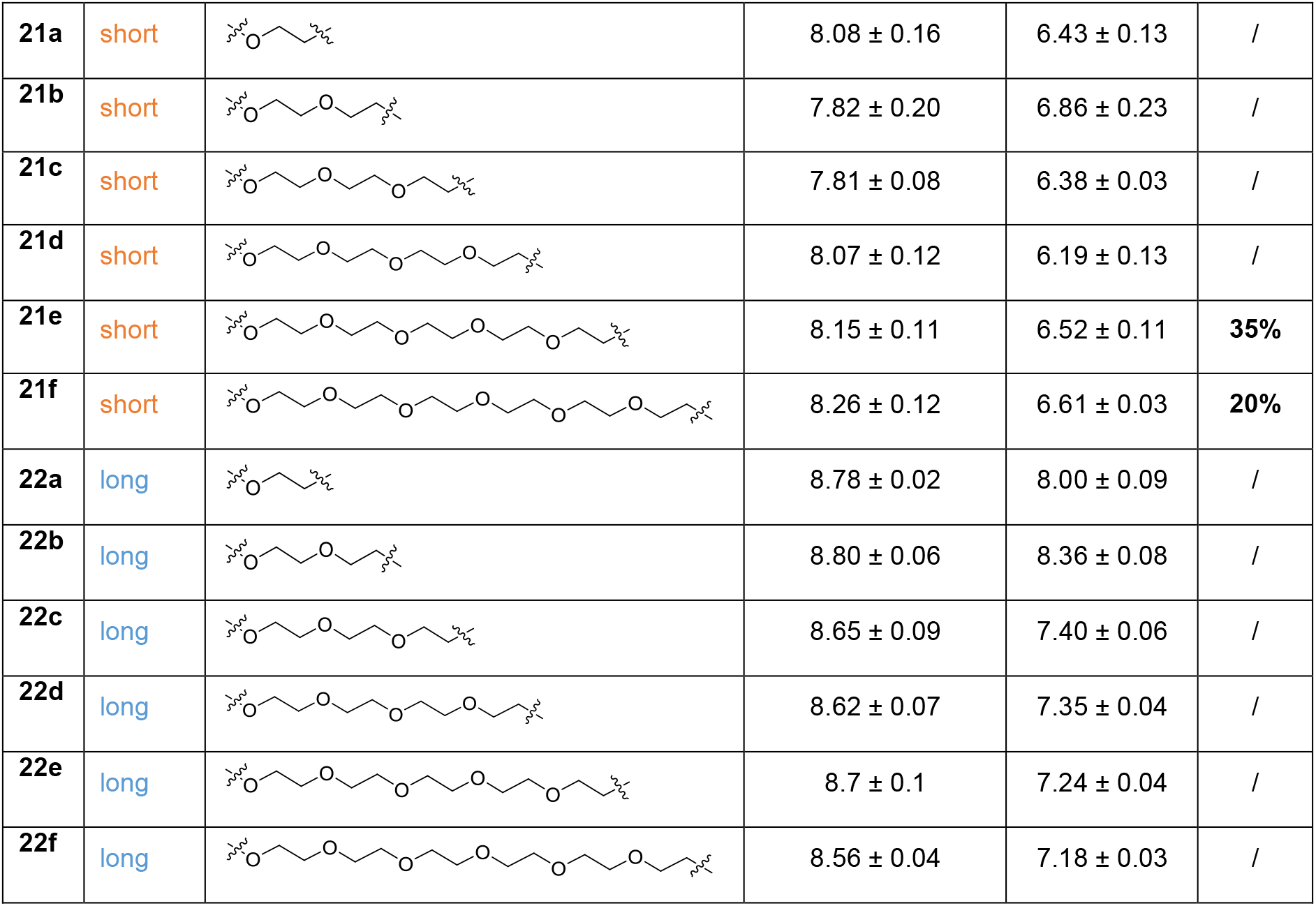
Synthesized sEH PROTACs with PEG linkers.

| no. | short/<br>long | linker | pIC<sub>50</sub>(hsEH) | pIC<sub>50</sub>(msEH) | D<sub>max</sub> |
| --- | --- | --- | --- | --- | --- |
| **21a** | short |  | 8.08 ± 0.16 | 6.43 ± 0.13 | / |
| **21b** | short |  | 7.82 ± 0.20 | 6.86 ± 0.23 | / |
| **21c** | short |  | 7.81 ± 0.08 | 6.38 ± 0.03 | / |
| **21d** | short |  | 8.07 ± 0.12 | 6.19 ± 0.13 | / |
| **21e** | short |  | 8.15 ± 0.11 | 6.52 ± 0.11 | **35%** |
| **21f** | short |  | 8.26 ± 0.12 | 6.61 ± 0.03 | **20%** |
| **22a** | long |  | 8.78 ± 0.02 | 8.00 ± 0.09 | / |
| **22b** | long |  | 8.80 ± 0.06 | 8.36 ± 0.08 | / |
| **22c** | long |  | 8.65 ± 0.09 | 7.40 ± 0.06 | / |
| **22d** | long |  | 8.62 ± 0.07 | 7.35 ± 0.04 | / |
| **22e** | long |  | 8.7 ± 0.1 | 7.24 ± 0.04 | / |
| **22f** | long |  | 8.56 ± 0.04 | 7.18 ± 0.03 | / |

In order to test potential PROTACs for degradation, Western Blots are predominantly used in the literature. As this method has a limited throughput and does not allow for an accurate determination of the PROTAC’s degradation efficiency (D_max_) and potency (DC_50_)^29^, a cellular NanoBiT based^22^ degradation assay for sEH was established in this study. The NanoBiT system allows to quantify the cellular level of a protein of interest (POI) by detection of luminescence generated by the small enzyme NanoBiT luciferase. In this setup, the POI is fused to a small peptide tag of 11 amino acids called HiBiT which represents a fragment of NanoBiT. HiBiT exhibits a very high affinity to the 18 kDa polypeptide LgBiT, the residual scaffold of luciferase. Complementation of the two fragments leads to a detectable bioluminescence signal. Thus, degradation of the sEH-HiBiT fusion protein is monitored by a decrease of bioluminescence (figure 4A). Due to its small size, HiBiT represents a minimally invasive tag and is suited for measuring protein levels in living cells. Furthermore, this assay setup allows for a precise calculation of the maximum percentage of degradation (D_max_), marking the PROTAC’s degradation efficiency, as well as the degradation potency DC_50_ (effective concentration to reach 50% of the PROTAC’s D_max_).

**Figure 4:**
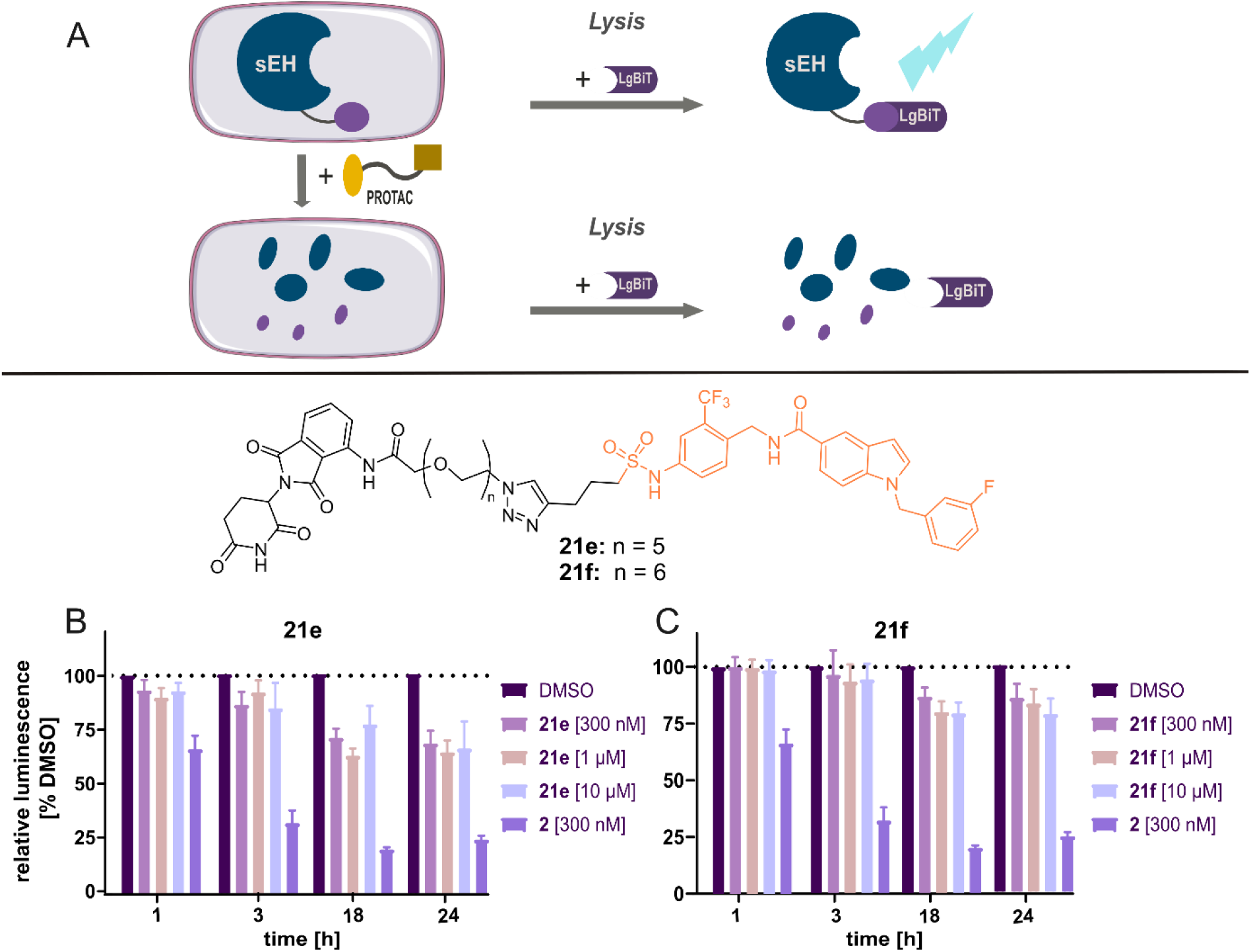
Development of a cellular NanoBiT based sEH degradation assay and testing of the synthesized compounds. A: NanoBiT assay principle. B and C: Results for short branch addressing sEH PROTACs **21e** with a PEG5 linker (Graph B) and **21f** PEG6 linker (Graph C). The best result within this series of PROTACs was obtained for **21e** with Dmax = 35%.

To obtain the respective cell line (HeLa^sEH-HiBiT^) the construct containing human sEH (aa1-aa555) followed by a linker and C-terminal HiBiT under the control of a constitutive HpGK promotor was used (for coding and protein sequences see SI). HeLa cells were stably transfected with the construct using the Sleeping Beauty method.^31^ The assay was set up in a 384 well plate format with 2000 cells per well in a total volume of 55 µL. After treatment of the cells with the respective compounds for different incubation intervals, cells are lysed and a LgBit and the NanoBiT are added (Promega Corporation), followed by luminescence detection using a plate reader. For the assay establishment, first-in-class sEH PROTAC **1a** was used as a positive control. PROTAC **2**^20^ was tested in eight concentrations from 0.001 µM to 3 µM along with DMSO treatment and cells were incubated for 1 h, 3 h, 18 h and 24 h, respectively. The obtained luminescence signals were plotted relative to DMSO against the concentration of **2** (figure sxy). In line with the previously published results we could not observe a total degradation of sEH. This phenomenon was explained by Wang et al. as they showed a selective degradation of cytosolic sEH while peroxisomal sEH was not degraded. Also in line with the literature, the hook effect was observed for a 3 µM concentration of **2** after 1 h and 3 h treatment. This commonly known PROTAC effect refers to less observed degradation with too high concentrations of PROTAC, leading to a preference of binary complex formations over ternary complex formation. Two control experiments were performed in order to validate the PROTAC induced degradation. First, cells were co-treated with **2** (300 nM) and sEH inhibitor GSK2256294 (**1**) (3 µM). As expected, this led to a rescue from degradation, indicating the degradation to be mediated by the PROTAC binding to the active site of sEH-H. Furthermore, cells were co-treated with **2** (300 nM) and the proteasome inhibitor MG132 (3 µM). We observed a restoration of the luminescence signal to nearly 100% DMSO, indicating a proteasomal degradation (figure Sxy). This finding is in contrast to the results of Wang et al., who had observed a rescue from degradation by the lysosome inhibitor Bafylomicin A1 (BafA1), while co-treatment with MG132 did not lead to a restoration of the sEH level in their setup.

In a first screening, we tested the synthesized sets of potential PROTACs in three concentrations (10.0, 1.0 and 0.1 µM) and incubated the HeLa^sEH-HiBiT^ cells for 1 h, 3 h, 18 h and 24 h. All PROTACs harbouring an alkyne linker (**18a-e** and **19a-e**) did not induce a decrease of luminescence compared to treatment with DMSO at any concentration or incubation time. The same was observed for long branch addressing compounds with PEG1-PEG6 (**22a-22f**) as well as short branch addressing compounds with PEG1-PEG4 linkers (**21a-21d**) (representative data shown in figure Sxy). However, the short branch addressing PROTACs **21e** and **21f**, exhibiting a PEG5 and PEG6 linker, induced a decrease of signal of 35% and 20% respectively (figure 4B and 4C). Thus, the most promising result within this series of PROTACs was obtained for **21e**, the combination of the short branch addressing inhibitor scaffold **4** and a PEG5 linker.

Based on these results, we decided to keep the short branch addressing inhibitor scaffold and the linker length of **21e** and change the CRBN ligand from the amide functionalized pomalidomide derivative to an ether functionalized thalidomide derivative. Structurally, the resulting compound **23** is a constitutional isomer of hit compound **21e**. Additionally, we aimed to examine whether the triazole-bridge of **21e** could have a disturbing effect on sEH degradation, and synthesized also the amide analogue **24** keeping the ether functionalized thalidomide on the CRBN recruiter side (scheme 5).

**Scheme 5:**
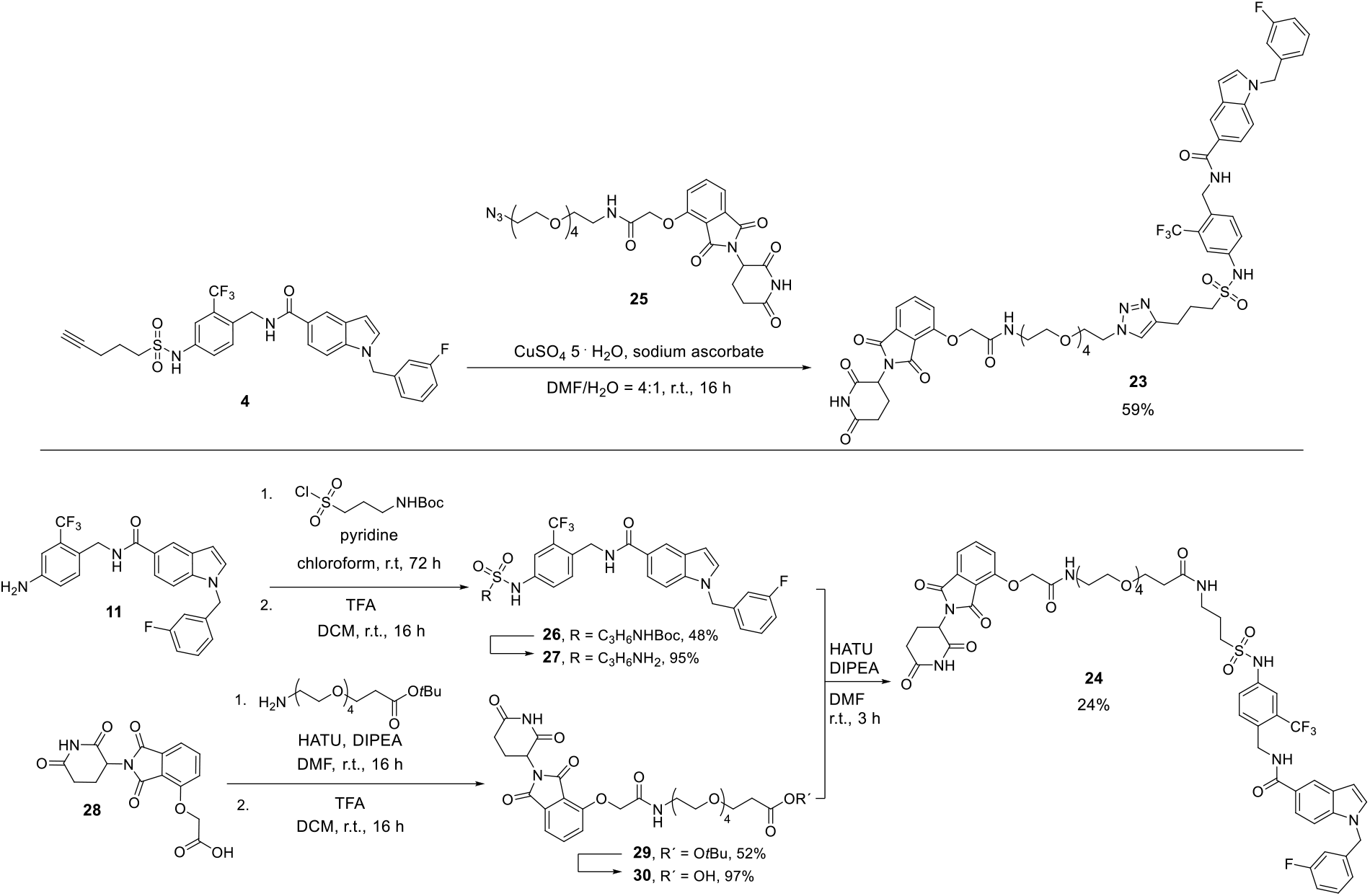
Synthetic routes for the synthesis of sEH PROTACs 23 and 24.

**23** was synthesized from short branch inhibitor scaffold **4** and the commercially available building block **25** under CuAAC conditions (scheme 5, upper part). In order to synthesize amide based PROTAC **24**, a new inhibitor scaffold **27** was synthesized with a primary amine handle for amide coupling. **27** was synthesized in two steps from previously synthesized aniline derivative **11**: A nucleophilic substitution reaction was performed with *tert*-butyl (3-(chlorosulfonyl)propyl)carbamate and pyridine to afford compound **26** which was Boc deprotection with TFA. Inhibitor scaffold **27** was then treated in an amide coupling reaction with building block **30** which had been synthesized in two steps before from **28** (scheme 5, lower part).

Both compounds were analyzed regarding their inhibitory potency towards hsEH-H and msEH-H as described for the first set of PROTACs (table 3). As observed before for other short branch addressing compounds, both PROTACs **23** and **24** exhibited excellent low nanomolar potencies towards hsEH-H, while up to two magnitudes lower potencies were observed towards the murine isoform. Interestingly, the triazole exhibiting PROTAC **23** was twice more potent than its amide containing analogue **24** towards hsEH-H and even more than thrice more potent towards msEH-H.

**Table 3:** Biochemical characterization of the optimized PROTACs 23 and 24 and the corresponding methylated PROTACs 31 and 32.

| no. | X | Y | pIC<sub>50</sub>(hSEH-H) | pIC<sub>50</sub>(mSEH-H) | pIC<sub>50</sub>(sEH-P) | pDC<sub>50</sub> | D<sub>max</sub> |
| --- | --- | --- | --- | --- | --- | --- | --- |
| **23** |  | H | 8.37 ± 0.04 | 7.12 ± 0.01 | inactive | 7.59 ± 0.12 | 75 % |
| **24** |  | H | 8.10 ± 0.07 | 6.60 ± 0.02 | inactive | 7.09 ± 0.06 | 80 % |
| **31** |  | CH<sub>3</sub> | 8.21 ± 0.08 | 6.67 ± 0.09 | n.d. | / | / |
| **32** |  | CH<sub>3</sub> | 7.95 ± 0.06 | 6.42 ± 0.07 | n.d. | / | / |

Next, both PROTACs were tested in the sEH-HiBiT-assay setup. In a first experiment, three concentrations were screened (0.3 µM, 1.0 µM and 3.0 µM) and cells were incubated for 1 h, 3 h, 18 h and 24 h. Already after a 1 h treatment partial degradation of 25 % was observed for triazole based PROTAC **23** (0.3 µM and 1.0 µM) and even a 50 % degradation for amide based PROTAC **24**. After 18 h, treatment resulted in a strong luminescence decrease of 75 % for **23** and 80% for **24**. Encouraged by these results, both compounds were subsequently tested in seven concentrations from 0.003 µM to 3.0 µM and with six different incubation times of 1 h-24 h. From these data, time dependent dose-response-curves were obtained (graphs 5A and 5B). Both PROTACs induced similarly strong degradation with D_max_ = 75 % and D_max_ = 80 %, respectively. In addition, both compounds exhibited high degradation potencies in the nanomolar range. Both compounds induced their highest D_max_ after 18 h while being slightly less effective after a 24 h treatment. The same has been observed before for the positive control **2** (scheme Sxy). To examine the effect of longer incubation, an additional experiment with incubation intervals of 18 h and 48 h was conducted for both compounds (300 nM). Indeed, less degradation was observed after a 48 h treatment with only 25% for PROTAC **23** and 30% for **24** (scheme Sxy). This finding may indicate a rapid re-synthesis of sEH in the cells, but could also be caused by a fast metabolism of the respective PROTAC in the cellular environment.

A set of control experiments^32^ was performed with both compounds **23** and **24** in order to validate the mechanism of action. In figure 5C representative results are shown for the triazole based PROTAC **23** (results for amide based PROTAC **24** are shown in figure Sxy). First, the corresponding negative control compounds **31** and **31** were synthesized (for experimental details see SI). These compounds are methylated at the glutarimide-N of the PROTAC’s thalidomide moiety and therefore should not bind to CRBN. No degradation is expected to be induced while exhibiting equivalent inhibitory potencies towards sEH-H as the respective PROTACs. Indeed, compounds **31** and **32** were strong sEH-H binders, but did not induce degradation (table 3 and figure 5C). Next, cells were co-treated with the respective PROTAC [300 nM] and an excess of the potent sEH-inhibitor GSK2256294A (**1**) or the CRBN binder pomalidomide [3 µM each]. Both co-treatments resulted in the restoration of the luminescence, indicating ternary complex formation to be essential for degradation. Also, with this experiment the compounds were shown to function via binding to the active site of sEH-H and not the sEH-P domain. This was additionally confirmed in an in vitro sEH-P activity assay, where PROTACs **23** and **24** resulted to be inactive (table 3). In order to test for proteasomal vs. lysosomal degradation, cells were co-treated with PROTAC **23** and the proteasome inhibitor MG132 or the lysosome inhibitor Bafylomicin A1 (Baf A1) respectively. As observed before for the positive control PROTAC **2**, co-treatment with MG132 resulted in signal restoration, while BafA1 did not affect degradation. Taken together, these results suggest the induced sEH degradation to be CRBN dependent and mediated by the proteasome.

**Figure 5:**
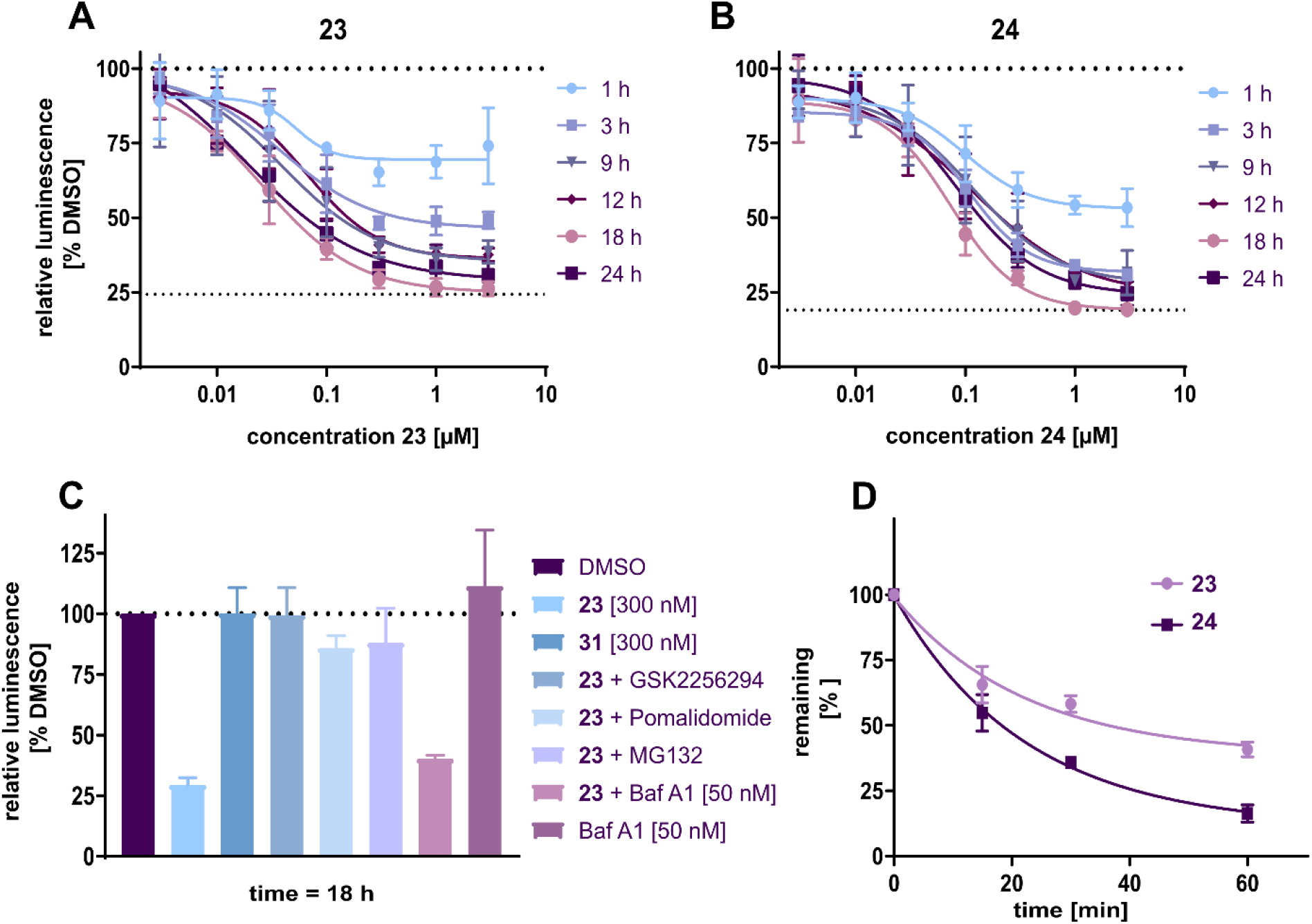
Biochemical characterization of optimized PROTACs **23** and **24**. A: degradation curves of triazole based PROTAC **23** for different incubation times (N=3). B: degradation curves of amide based PROTAC **24** for different incubation times (N=3). C: Control experiments for PROTAC 23 (N=3). D: Evaluation of metabolic stability of PROTACs **23** and **24** in rat liver microsomes.

Optimized PROTACs **23** and **24** were further characterized regarding their pharmacokinetic properties. In a simple solubility test the water solubility of both PROTACs was determined to be in a range between 3 µM and 5 µM (measured in DPBS buffer, 0.5 % DMSO, results shown in figure Sxy). Furthermore, both compounds were tested for cytotoxicity in a Cell Titer Glo assay in HepG2 cells. After 72 h, the live rate was still at about 100% even for the highest tested concentration (50 µM), proving the compounds not to be cytotoxic in the concentration range they are used (figure Sxy). Lastly, the metabolic stability was determined in both murine and rat liver microsomes. In rat liver microsomes, only 16% of **24** remained after 60 min, whereas **23** remained with 41% (figure 5D). Both species were more stable in mouse liver microsomes, with 48% remaining **23** and 34% remaining **24**, but also here **23** resulted to be more stable (figure Sxy). Based on these results and the higher degradation potency (table 3), triazole containing PROTAC **23** was chosen and tested in further cell based experiments.

In order to examine the applicability of PROTAC **23** as a tool compound in cell based systems, further experiments were performed in primary human cells. Immunofluorescence staining experiments were performed in human M1 macrophages (figure 6).

**Figure 6:**
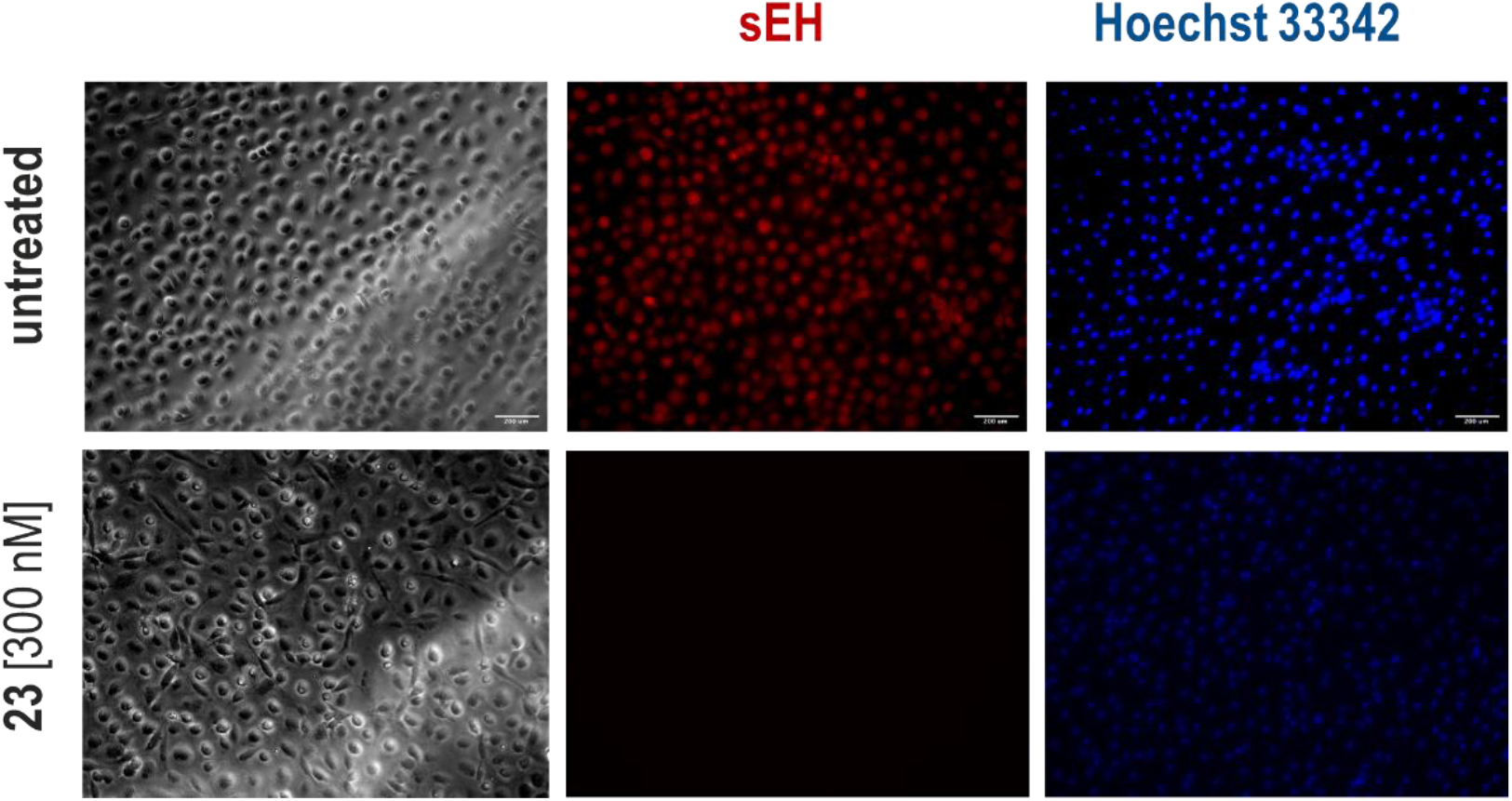
Immunofluorescence staining experiments of sEH in human primary M1 macrophages treated with 300 nM **23** or untreated (DMSO). Cell nuclei and sEH were visualized by fluorescence microscopy at 10x magnification. Representative experiment is shown. Scale bar: 200 μm. n=3.

The cells were treated with 300 nM PROTAC **23** or 0.5% DMSO and incubated for 18 h. Afterwards, cells were incubated with a rabbit anti-human sEH antibody, followed by incubation with an Alexa Fluor 633 conjugated secondary antibody (goat anti-rabbit) for 1 h. Cell nuclei were marked with Hoechst 33342. Immunofluorescence staining of sEH (λ_ex_: 590-650 nm, λ_em_: 662-738 nm) and cell nuclei (λ_ex_: 325-375 nm, λ_em_: 435-485 nm) were detected using a Leica fluorescence microscope. Figure 6 shows the contrast between untreated (upper panel) and treated cells (lower panel): while sEH could be well detected before treatment, there was a strongly reduced sEH signal of 99% after treatment with 300 nM, but cell nuclei remained detectable. This result suggests efficient sEH degradation induced by PROTAC **23**.

The obtained results proof the applicability of PROTAC **23** to study the function of sEH in primary cells, marking the successful translation to non-artificial systems.

## Conclusion

In conclusion, we have developed the potent sEH PROTAC **23** based on the previously published sEH inhibitor FL217. The rational design of PROTAC **23** was based on the crystal structure of previously published sEH inhibitor **3** and a structure-activity-relationship (SAR) investigation was performed with a total of 24 synthesized sEH PROTACs. Furthermore, a cell-based sEH degradation assay was developed based on the HiBiT-technology in order to allow a rapid evaluation of the synthesized compounds. PROTAC **23** was identified to be a highly potent degrader and exhibited acceptable solubility and metabolic stabilit for the application in cell based biological systems. **23** was successfully applied to degrade sEH in primary human, demonstrating its applicability as a tool compound to study sEH in non-artificial cell based systems.

